# Neonatal multi-modal cortical profiles predict 18-month developmental outcomes

**DOI:** 10.1101/2021.09.23.461464

**Authors:** Daphna Fenchel, Ralica Dimitrova, Emma C. Robinson, Dafnis Batalle, Andrew Chew, Shona Falconer, Vanessa Kyriakopoulou, Chiara Nosarti, Jana Hutter, Daan Christiaens, Maximilian Pietsch, Jakki Brandon, Emer J. Hughes, Joanna Allsop, Camilla O’Keeffe, Anthony N. Price, Lucilio Cordero-Grande, Andreas Schuh, Antonios Makropoulos, Jonathan Passerat-Palmbach, Jelena Bozek, Daniel Rueckert, Joseph V. Hajnal, Grainne McAlonan, A. David Edwards, Jonathan O’Muircheartaigh

## Abstract

Developmental delays in infanthood often persist, turning into life-long difficulties, and coming at great cost for the individual and community. By examining the developing brain and its relation to developmental outcomes we can start to elucidate how the emergence of brain circuits is manifested in variability of infant motor, cognitive and behavioural capacities. In this study, we examined if cortical structural covariance at birth, indexing coordinated development, is related to later infant behaviour. We included 193 healthy term-born infants from the Developing Human Connectome Project (dHCP). An individual cortical connectivity matrix derived from morphological and microstructural features was computed for each subject (morphometric similarity networks, MSNs) and was used as input for the prediction of behavioural scores at 18 months using Connectome-Based Predictive Modeling (CPM). Neonatal MSNs successfully predicted social-emotional performance. Predictive edges were distributed between and within known functional cortical divisions with a specific important role for primary and posterior cortical regions. These results reveal that multi-modal neonatal cortical profiles showing coordinated maturation are related to developmental outcomes and that network organization at birth provides an early infrastructure for future functional skills.

## 1. Introduction

Developmental delays occur in around 13% of infants in the US population (Rosenberg et al. 2008). Delays can be observed in motor, cognitive, language and communicative domains, and when they persist, they are termed developmental disabilities. These can place a great emotional and financial burden on the individual, their family, and the general community (Shahat & Greco, 2021; Stabile & Allin, 2012). While some infants catch up with their peers, others will continue to present difficulties (Riva et al. 2021). Behavioural delays are associated with a higher likelihood of autism spectrum conditions (ASC), attention deficit hyperactivity disorder (ADHD) and schizophrenia (Gurevitz et al. 2014; Landa & Garrett-Mayer, 2006; Sorensen et al. 2010). Within the general population, developmental milestones are related to cognitive functions both in childhood and adulthood (Flensborg-Madsen & Mortensen, 2018; Murray et al. 2007), reiterating their importance over the lifespan.

Risk factors for poor neurodevelopmental outcomes include familial history of neurodevelopmental conditions (Ozonoff et al. 2011; Shivers et al. 2019; Stromswold, 1998) and inherited or *de novo* genetic changes (Cooper et al. 2011; Marshall et al. 2008), in addition to preterm birth, low birth weight (Aylward, 2014; Pascal et al. 2018) and other perinatal complications (Mwaniki et al. 2012). However, most infants will have no recognized predisposing factor. Adding to this complexity, the pace of motor and language development in the first years of life is variable within and between individuals even within the ‘normal’ range (Fenson et al. 1994; Piek, 2002). Identifying individuals at potentially greater likelihood for difficulties allows for early interventions which have been found to improve outcome (Dawson et al. 2010; Jeong et al. 2021). Specifically, by being able to recognize babies in the general population who might need extra support, we can begin to address difficulties or potential difficulties very early on, while brain development is still in its early sensitive period (Reh et al. 2020).

Prospectively profiling the developing brain and investigating its relationship with adaptive and maladaptive behaviours, promotes our understanding of innate and external factors contributing to variability, vulnerability, and resilience to adverse outcomes. MRI studies have concluded that structural and functional brain networks start to develop in the fetal period and continue to fine-tune during childhood (Batalle et al. 2018). However how (and when) this emerging brain architecture relates to behavioural outcomes in infanthood is yet to be determined. The majority of early developmental studies have focused on brain-behaviour relationships in preterm neonates, a group that on average has a greater likelihood of delayed or atypical development (Van’t Hooft et al. 2015). More recently, a conceptual shift attempts to move from association to prediction (Rosenberg et al. 2018), with more studies examining brain structure in the term-born neonatal population with no apparent risks for poorer developmental outcomes (Girault et al. 2019b; Wee et al. 2017). Most attention has been given to the predictive ability of white matter connectivity (Ball et al. 2015; Girault et al.2019b; Keunen et al. 2017; Wee et al. 2017).

Morphometric similarity networks (MSNs) (Seidlitz et al. 2018) are based on structural covariance between brain regions whereby similarity is thought to reflect synchronized maturation and relatedness (Alexander-Bloch et al. 2013a, 2013b). The origins of this coordinated development may be the result of sharing an early progenitor, or from exposure to similar early signalling, in a process that can be modulated by genetic and environmental exposures (Alexander-Bloch et al. 2013a). Accordingly, related regions likely reflect a joint functional purpose, with a similar transcriptomic profile (Yee et al. 2018). It has been hypothesized that abnormal patterns of brain covariance may result from atypicality in the establishment of the first connections innervating the cortical plate, starting at mid-gestation (Bullmore et al. 1998). This has implications for efficient information transfer and functionality and therefore could potentially serve as an early marker for the development of mental and motor abilities.

MSNs incorporate multiple MRI modalities into the estimated structural covariance, both microstructural and morphological, to overcome the limitations of using individual features with particular spatio-temporal trajectories. These provide a more comprehensive description of the brain, improving the predictive ability of clinical symptoms and behaviour from brain data (Liu et al. 2015; Tulay et al. 2019). Correspondingly, covariance networks based on multiple MRI measures are better at capturing the underlying cellular composition compared to structural covariance networks based on a single measure (Seidlitz et al. 2018). In adults, MSNs are related to cognitive abilities and the expression of genes associated with neurodevelopmental conditions (Morgan et al. 2019; Seidlitz et al. 2018,2020).

In our previous work, we used this method to characterize the developing brain at the neonatal timepoint using structural and diffusion indices (Fenchel et al. 2020), reporting a community structure largely aligned with known functional distinctions and network temporal trajectories, and showing close similarity with cytoarchitectural features (Ball et al. 2020). In this current study, we were interested in furthering our understanding of how this neonatal cortical organization relates to infant developmental outcomes. Therefore, here we asked whether cortical profiles at term-birth, derived from MSNs, are associated with- and predictive of- motor, cognitive, language and social-emotional abilities at 18 months. We attempted to predict infant behaviour from neonatal MSNs using connectome-based predictive modelling (CPM), a data-driven linear approach to predict continuous measures of behaviours from individual connectivity matrices (Shen et al. 2017). Following CPM, we examined if network-strength summary measures at the whole cortex level and within cortical functional clusters were able to capture the same brain-behaviour patterns observed at the single-edges level.

## 2. Methods

### 2.1 Subjects

This study included a sample of term-born healthy neonates participating in the Developing Human Connectome Project (dHCP); (http://www.developingconnectome.org/), scanned at the Newborn Imaging Centre at Evelina London Children’s Hospital, London, UK. Images are openly available on the project website. This project has received ethical approval (14/LO/1169) and written informed consent was obtained from parents. As part of the dHCP project, subjects are invited for a follow-up visit to assess infant development at 18 months. This assessment includes The Bayley Scales of Infant and Toddler Development (Bayley-III) (BSID) (Bayley, 2006) exploring overall developmental aspects, as well as the Quantitative Checklist for Autism in Toddlers (Q-CHAT) (Allison et al., 2008) for assessment of social-emotional development. Out of the 241 subjects included in the initial analysis and for which MSNs were constructed (Fenchel et al., 2020), n=204 completed the Bayley-III assessment and n=198 completed the Q-CHAT assessment. Only subjects with information on a proxy of socio-economic status, the Index of Multiple Deprivation (IMD) (https://tools.npeu.ox.ac.uk/imd/), were included to control for its possible confounding effect. This resulted in a sample size of n=193 with Bayley-III data and n=187 with Q-CHAT data.

### 2.2 Image acquisition and processing

Neonatal MR brain images were acquired on a 3T Philips Achieva scanner without sedation, using a dedicated 32-channels head coil system (Hughes et al., 2017). Acquisition, reconstruction and processing of structural and diffusion images followed optimized protocols for the neonatal brain implemented as part of the dHCP pipeline and have been previously described in Fenchel et al. 2020. T2-weighted (T2w) images were obtained using a turbo spin-echo (TSE) sequence, acquired in sagittal and axial planes with TR=12s, TE=156ms, SENSE factor 2.11 (axial) and 2.58 (sagittal) with overlapping slices (resolution 0.8×0.8×1.6mm). T1-weighted (T1w) images were acquired using an Inversion Recovery TSE sequence with the same resolution using TR=4.8s, TE=8.7ms, SENSE factor 2.26 (axial) and 2.66 (sagittal). Structural images were reconstructed to a final resolution of 0.5×0.5×0.5mm, using slice-to-volume registration (Cordero-Grande et al. 2018). Structural processing followed the pipeline described in (Makropoulos et al., 2018): Motion- and bias-corrected T2w images were brain extracted and segmented. White, pial and midthickness surfaces were generated, inflated and projected onto a sphere. Brains were aligned to the 40-week dHCP surface template (Bozek et al., 2018) using Multimodal Surface Matching (MSM) (Robinson et al. 2013, 2014). Cortical features including cortical thickness (CT), pial surface area (SA), mean curvature (MC), and the T1w/T2w ratio indicative of myelin content (MI) were extracted for each subject (Makropoulos et al., 2018).

Diffusion images were obtained using parameters TR=3.8s, TE=90ms, SENSE factor=1.2, multiband=4, partial Fourier factor=0.86, resolution 1.5×1.5×3.0mm with 1.5mm overlap (Hutter et al. 2018). Diffusion gradient encoding included images collected at b=0s/mm^2^ (20 repeats), b=400s/mm^2^ (64 directions), b=1000s/mm^2^ (88 directions), b=2600s/mm^2^ (128 directions) (Tournier et al. 2020). Diffusion images were denoised (Veraart et al. 2016), Gibbs-ringing suppressed (Kellner et al. 2016), and the field map was estimated (Andersson et al. 2003). Images were corrected for subject motion and image distortion with slice-to-volume reconstruction using multi-shell spherical harmonics and radial decomposition (SHARD) and were reconstructed to a final resolution of 1.5×1.5×1.5mm (Christiaens et al. 2021). A tensor model was fitted using a single shell (b=1000s/mm^2^), and fractional anisotropy (FA) and mean diffusivity (MD) maps were generated using MRtrix3 (Tournier et al. 2019). Neurite density index (NDI) and orientation dispersion index (ODI) maps were calculated using the default NODDI toolbox implementation with default values (Zhang et al. 2012). Diffusion maps were registered onto individual T2w images using FSL’s epi_reg (FLIRT) and then projected onto the cortical surface using Connectome Workbench. All images were visually inspected for motion or image artefacts and data excluded accordingly (Fenchel et al. 2020), and images were checked for registration errors.

### 2.3 MSNs construction

MSN construction for this cohort was described previously in Fenchel et al. 2020 and is summarized in Figure 1. Briefly, the cortical surface was parcellated into 75 bilateral equal-sized regions with Voronoi decomposition. Seven of these regions were excluded due to diffusion signal dropout. Each region was then characterized by an eight-feature vector of mean normalized values of four structural features: CT, MC, MI and SA and four diffusion features: FA, MD, NDI and ODI. Pearson’s correlation between the eight-feature vector for every pair of regions was calculated, resulting in a 143 x 143 similarity-based connectivity matrix for each subject. Values were Fisher’s-z-transformed before analysis.

**Figure 1.**
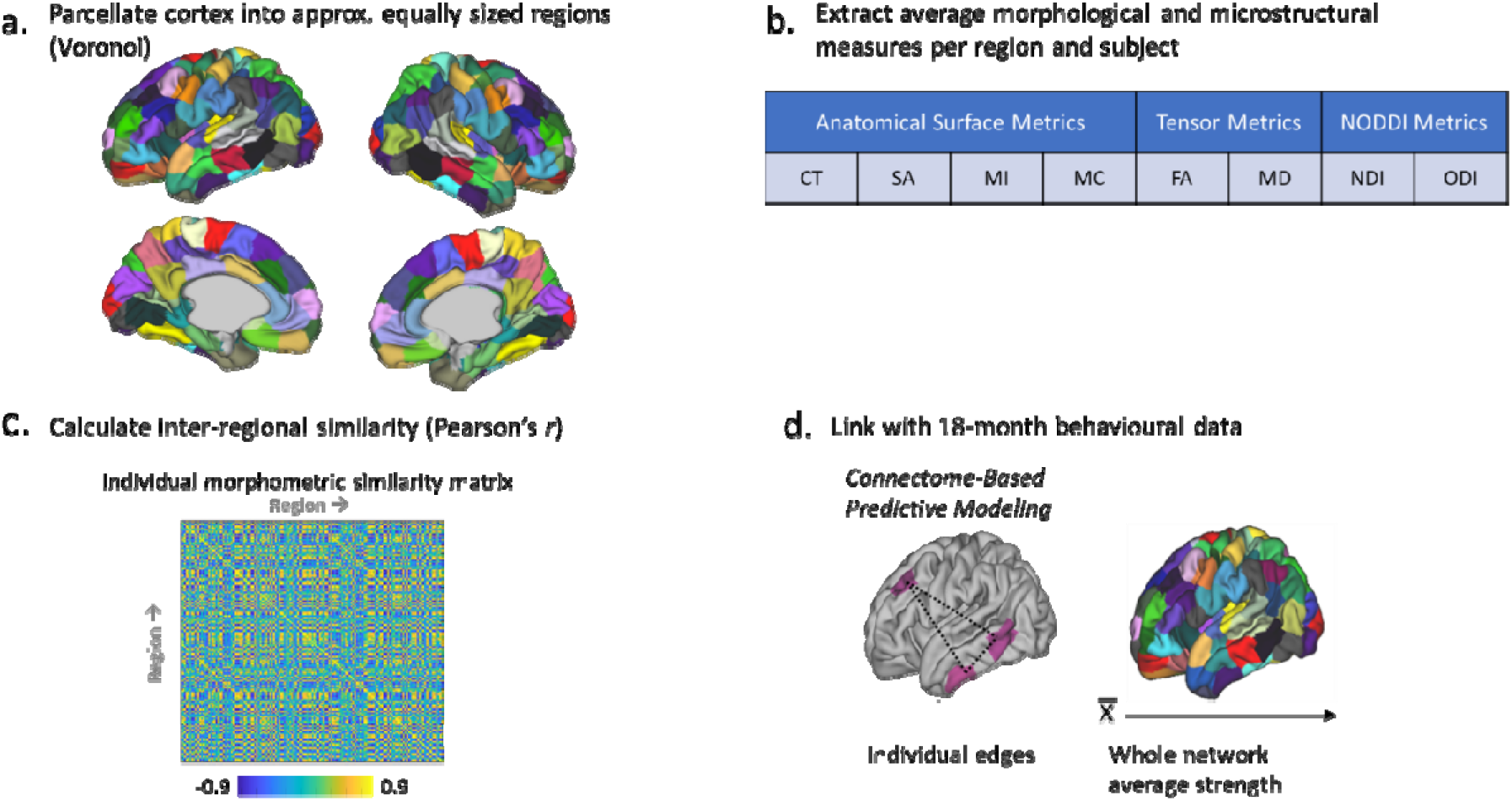
Pipeline Morphometric Similarity Networks construction and behavioural analysis. a. Regions are defined using Voronoi tessellation of the cortical surface; b. A feature vector of averaged normalized values of cortical thickness (CT), mean curvature (MC), myelin index (MI), surface area (SA), fractional anisotropy (FA), mean diffusivity (MD), neurite density index (NDI) and orientation dispersion index (ODI) is derived for each region; c. Each pair of regions is correlated using Pearson’s r, resulting in an individual similarity-based connectivity matrix. d. Network strength at a whole-network level and single-edges level is related to behavioural measures by means of association and prediction respectively.

### 2.4 Behavioural developmental assessment

Mean age at developmental assessment was 18.69±1.04 months (range 17.16-24.46 months), mean corrected age for gestational age (GA) at birth was 18.68±1.00 months (range 17.30-24.33 months), the latter used for Bayley-III score calculation. The Bayley-III (Bayley, 2006) is a commonly used tool for tracking infants’ development, targeting to identify possible developmental delays. Standardized scores are divided into a motor composite score, derived from gross and fine motor scaled sub-scales, a language composite score derived from expressive and receptive language scaled sub-scales and a cognitive composite score. Bayley-III assessments were completed by trained practitioners. A higher score on these scales reflects better performance.

Social-emotional development was determined by the Q-CHAT (Allison et al. 2008), a parent-based report of 25 items including joint attention, pretend play, language development, repetitive behaviours, and social communication. A higher summary score suggests more social-emotional difficulties, possibly indicating early autistic traits

### 2.5 Prediction of developmental outcomes from MSNs

We utilized Connectome-based Predictive Modeling (CPM) (Shen et al. 2017) to explore the predictive ability of neonatal MSNs for four developmental outcomes measures, the three Bayley-III composite scores (cognitive, language and motor) and the Q-CHAT score. The model is trained each time on n-1 subjects and is tested on the left-out subject for calculation of the predicted behavioural scores (leave-one-out cross-validation, LOOCV). Each individual edge was partially correlated (Spearman’s r) with the behavioural measure, controlling for postmenstrual age (PMA) at scan, sex, total intracranial volume (ICV), IMD and time from birth to scan. Positive edges (associated with higher behavioural scores) and negative edges (associated with lower behavioural scores) with p<0.05 were then selected. For each subject in the training set, these edges are summed separately to create a positive network sum and a negative network sum. Linear regression with no intercept (Rosenberg et al. 2018; Shen et al. 2017) linking the positive and negative sums was then performed. The predicted behaviour for the left-out subject is calculated by fitting this subject’s sum of positive and negative edges identified in the training set, adjusted for covariates, with the beta coefficients derived from the training model. Model performance was assessed by computing Spearman’s r between the observed and predicted behaviours and the root mean square error (RMSE). This performance was assessed by generating a null distribution of r values from N=999 random permutations of the behavioural data. The resulting p-value is calculated as the number of r values equal or larger to the original r value divided by N+1. Successful models using LOOCV were further examined for robustness using 10-fold cross-validation for 100 iterations. In each iteration, subjects are randomly assigned to each of 10 groups, where each time a different group serves as the test set and the remaining nine groups serve as the training set. Spearman’s r, associated p-value, and RMSE were calculated for each iteration and then averaged.

Predictive networks were determined as significant if passed both cross-validation methods. These were defined by taking edges appearing in at least 90% of testing runs in the LOOCV. For clarity and ease of interpretation, predictive edges were examined in the context of the seven clusters reported before in Fenchel et al. (2020): occipital & parietal, limbic, anterior frontal, insular & medial frontal, fronto-temporal, cingulate and somatosensory & auditory. For each cluster, we (1) summed separately the number of predictive positive and negative edges within the cluster and divided that by the number of all possible edges within that cluster to control for cluster size and (2) summed separately the number of positive and negative edges between each pair of clusters and divided that by the number of all possible edges between those pairs.

### 2.6 Association between MSNs summary measures and developmental measures

The association between whole-network average strength (across the entire cortex) and the eight developmental measures was examined by averaging a symmetric triangle of the connectivity matrix, excluding self-connections. This was entered together with PMA at scan, sex, ICV, IMD and time from birth to scan into a general linear model where the developmental measure was the dependent variable. Partial R^2^ for the brain network measure was calculated as (SSE reduced model-SSE full model)/SSE reduced model. Although PMA at scan, ICV and time from birth to scan were significantly correlated (r=0.71, p<0.001), no variance inflation factor (VIF) exceeded 5 (Craney & Surles, 2002) and therefore we retained all covariates for all analyses.

## 3. Results

### 3.1 Demographics and behavioural scores

Demographics of the sample and mean behavioural scores are presented in Table 1. All Bayley-III items were positively correlated with each other and negatively correlated with the Q-CHAT (Supplementary Table 1). Corrected age at assessment was not associated with either the Bayley-III scores or the Q-CHAT score.

**Table 1.** Demographics and behavioural scores.

|  |  | **N(%) / Median(range)** |
| --- | --- | --- |
| **Demographics** | Sex (male) | 101 (52.3%) |
|  | GA at birth (weeks) | 40.14 (37.29-42.14) |
|  | PMA at scan (weeks) | 40.86 (37.43-44.43) |
|  | Time from birth to scan (weeks) | 0.29 (0-5.28) |
|  | IMD | 26.12 (1.55-61.37) |
|  |  | **Mean±SD** |
| **Bayley-III** | Cognitive composite | 99.95±10.19 |
|  | Language composite | 96.39±15.41 |
|  | Expressive language | 8.81±2.58 |
|  | Receptive language | 9.90±3.14 |
|  | Motor composite | 101.41±9.70 |
|  | Fine motor | 11.39±2.20 |
|  | Gross motor | 9.02±1.92 |
| **Q-CHAT** |  | 30.65±9.19 |

GA- gestational age, PMA- postmenstrual age, IMD- Index of Multiple Deprivation, Bayley-III- Bayley Scales of Infant and Toddler Development, Q-CHAT- Quantitative Checklist for Autism in Toddlers

### 3.2 Single edges prediction-CPM

Neonatal MSNs successfully predicted the Q-CHAT score (*r*_s_=0.196, p=0.007, p permute=0.019, RMSE=9.53) and the language composite score (*r*_s_=0.182, p=0.011, p permute=0.024, RMSE=16.30) (Figure 2, Supplementary Figure 1). However, only the Q-CHAT network remained significant following 10-fold cross-validation (Q-CHAT *r*_s_ =0.188, p=0.015, RMSE=9.60; language *r*_s_ =0.116, p=0.163, RMSE=16.80) and was therefore retained as the only robust predictive model.

**Figure 2.**
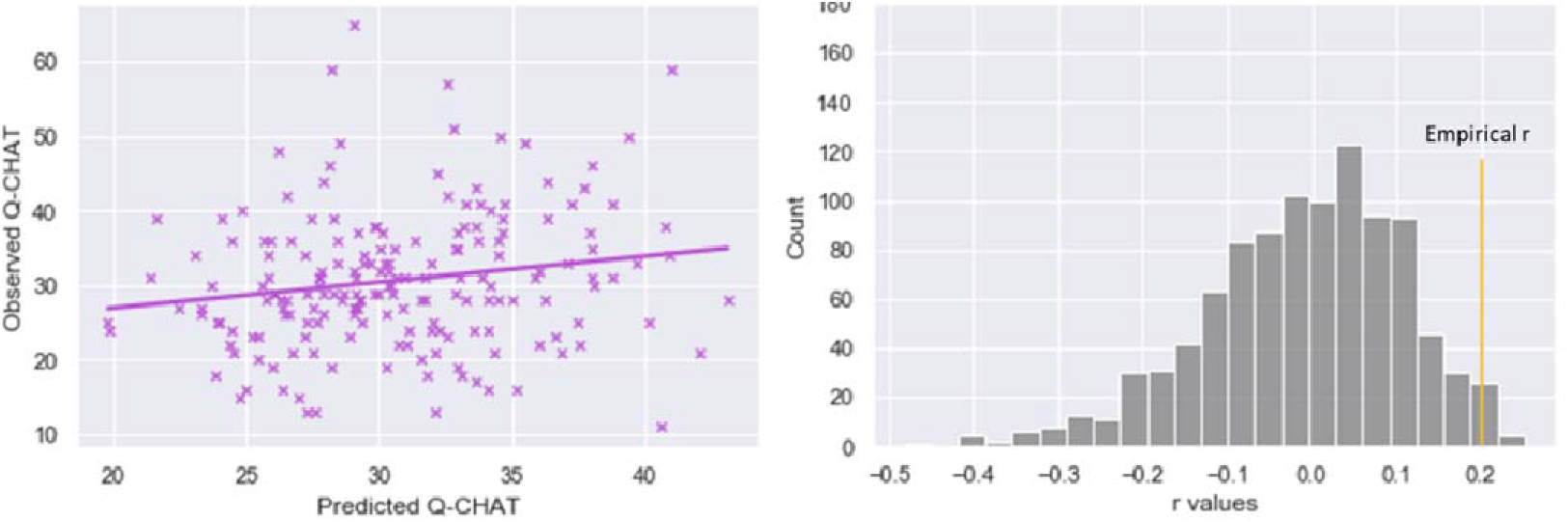
Prediction of Q-CHAT scores from neonatal MSNs using CPM-CPM. Plots of significant correlation between predicted and observed Q-CHAT scores (left) and results of null r values with permutation testing (right) using Connectome-based Predictive Modeling (CPM).

The positive and negative predictive networks for the Q-CHAT included 1.6% and 2.8% of all possible connections, respectively. The proportion of edges within each of the seven clusters included in the predictive networks is presented in Figure 3. For Q-CHAT, the highest proportion of positive predictive edges was within the anterior frontal and occipital and parietal clusters, while the cingulate did not show any within-cluster predictive edges. For the negative predictive network however, the highest proportion of edges was observed in the cingulate cluster. Limbic edges were not part of the negative or positive networks (Figure 3).

**Figure 3.**
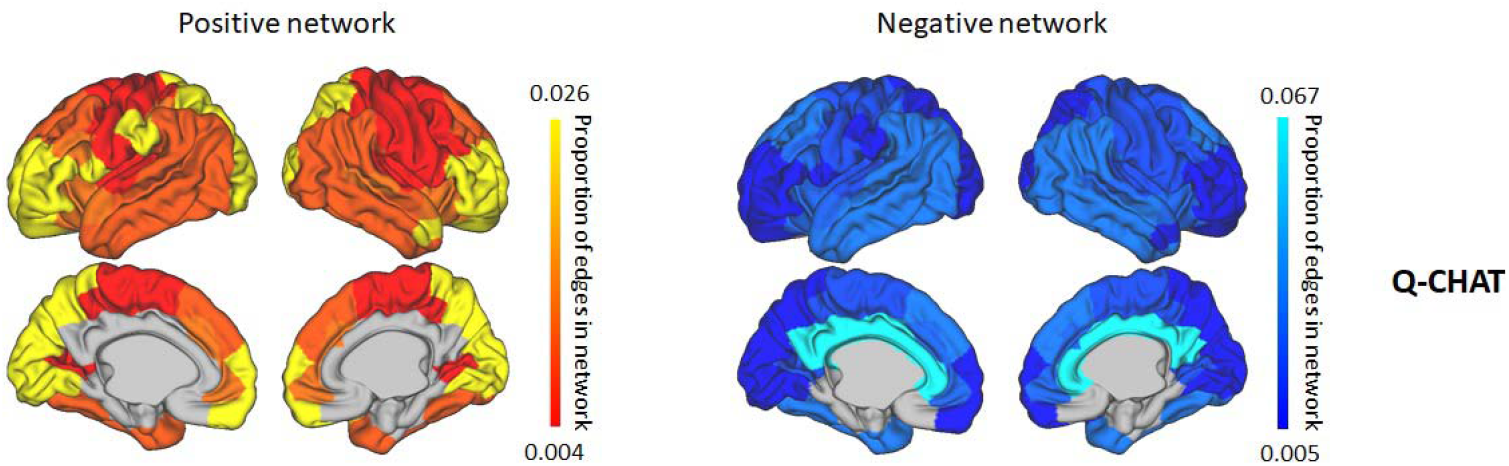
Proportion of within-cluster edges involved in social-emotional networks. Proportion of edges included in successful prediction model of social-emotional outcomes connecting nodes within each of the seven clusters.

The proportion of predictive edges connecting between clusters is presented in Figures 4. While the relationship emerging between clusters is complex, the highest proportion of predictive edges was observed for the occipital and parietal cluster. Interestingly, although we do not consider the language network as a robust enough predictor, the observed patterns of the language network seem to be closely related to the Q-CHAT network (Supplementary Figure 2): the positive network of the Q-CHAT was similar to the negative network of the language composite, and the negative network of the Q-CHAT was similar to the positive network of the language composite (Figure 4, Supplementary Figure 2). Therefore we looked at the overlap between these networks and found that 16.6% of edges in the negative language network overlapped with the positive Q-CHAT network and 26.8% of edges in the positive language network overlapped with the negative Q-CHAT network. No overlapping edges were found between the two positive networks or the two negative networks.

**Figure 4.**
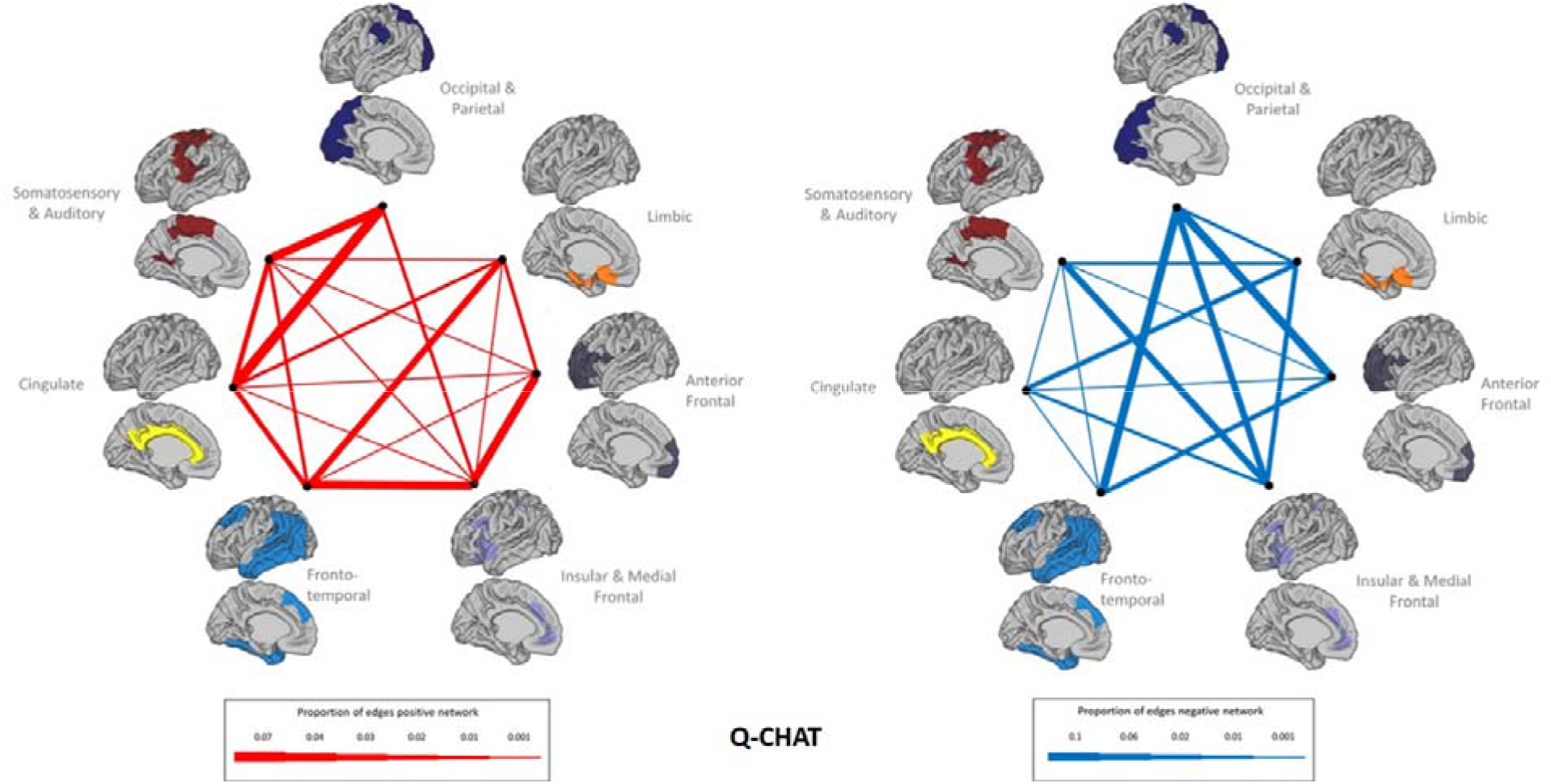
Proportion of between-cluster edges involved in social-emotional networks. Proportion of edges included in Q-CHAT prediction model connecting nodes between clusters. On the left the positive network is shown and, on the right, the negative network.

### 3.3 Whole-network average strength

There were significant positive associations between whole-network average strength and language composite and expressive language sub-scores, and a negative significant association with the Q-CHAT scores. No associations were found for the motor composite and associated sub-scales or the cognitive composite (Figure 5, Table 2). Variance explained by the full model was 7% for expressive language, 10% for language composite, and 14% for Q-CHAT, with specific contribution of network strength estimated at 4%, 3% and 4% respectively (partial R^2^) (Table 2).

**Figure 5.**
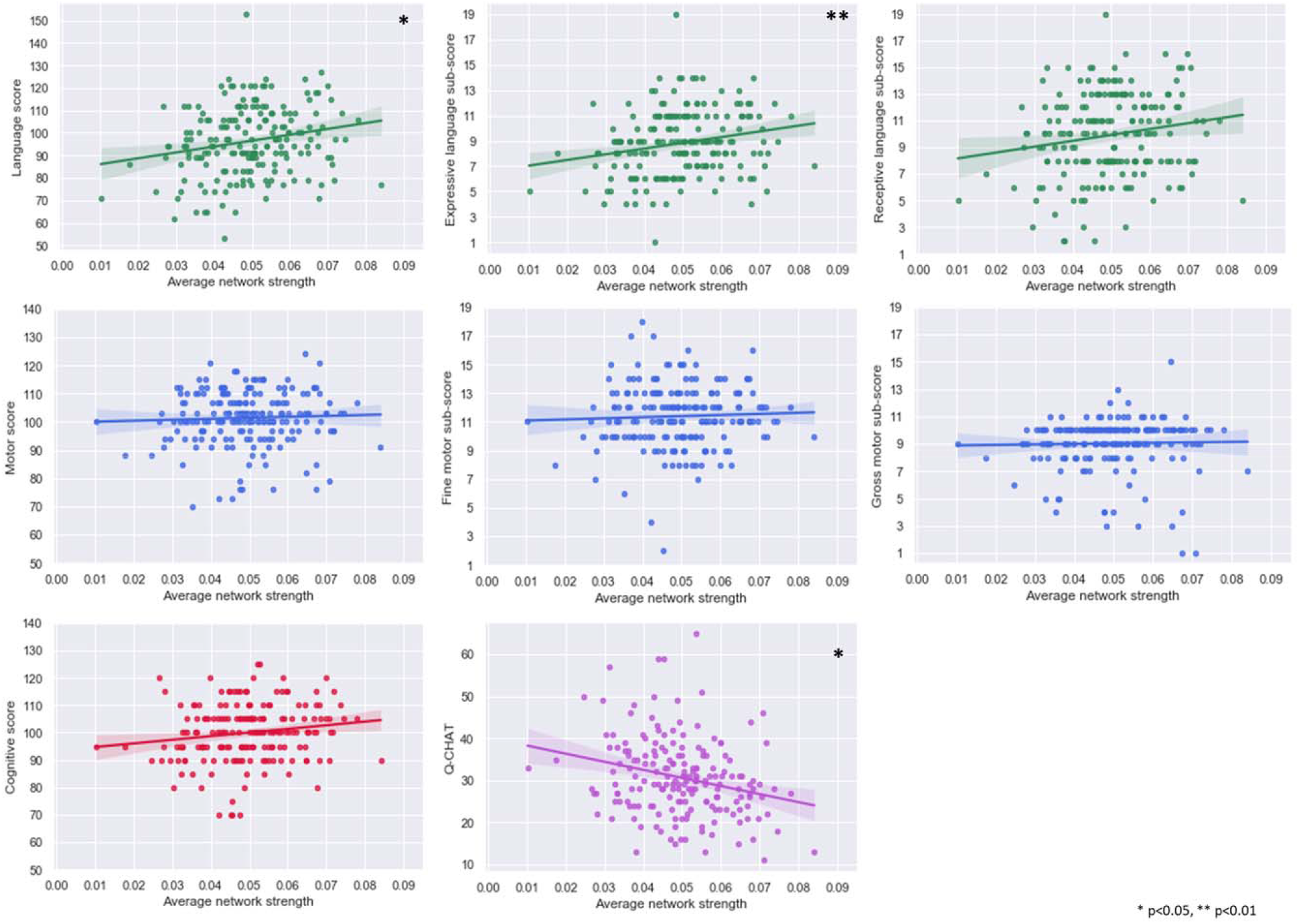
Scatterplots of whole-network average and behaviour. Plots of average MSN strength across the cortex against language, motor, cognitive and social-emotional measurements 18 months.

**Table 2.** Linear regression results for whole-network average strength and behaviour.

|  | Beta (95% CI) | SE | T-value | R<sup>2</sup> full ANOVA model |
| --- | --- | --- | --- | --- |
| Cognitive composite | 114.78 (-19.19-248.75) | 67.91 | 1.69 | 0.07 |
| Language composite | 221.91 (23.14-420.67) | 100.75 | 2.20 * | 0.10 |
| Expressive language | 45.58 (11.67-79.49) | 17.90 | 2.65 ** | 0.07 |
| Receptive language | 31.02 (-9.32-71.36) | 20.45 | 1.52 | 0.11 |
| Motor composite | 39.97 (-91.57-171.51) | 66.68 | 0.6 | 0.01 |
| Fine motor | 6.0 (-23.50-35.50) | 14.95 | 0.4 | 0.03 |
| Gross motor | 7.19 (-18.73-33.10) | 13.14 | 0.55 | 0.02 |
| Q-CHAT | -153.78 (-271.31-(-36.26)) | 59.56 | -2.58 * | 0.14 |

\* p<0.05, \*\* p<0.01; SE-Standard error, Q-CHAT- Quantitative Checklist for Autism in Toddlers.

To elucidate whether network strength is valuable in the context of the community structure (clusters) of neonatal MSNs reported previously (Fenchel et al. 2020), we examined the average network strength within each of the seven clusters. The network strength within the insular and medial frontal cluster was related to scores on the language composite (β=25.82, 95% CI=3.30-48.33, p<0.05, R^2^ full=0.11, R^2^ partial=0.03), receptive language sub-scale (β=5.37, 95% CI=0.84-9.91, p<0.05, R^2^ full=0.13, R^2^ partial=0.03), cognitive composite (β=15.44, 95% CI=0.30-30.58, p<0.05, R^2^ full =0.08, R^2^ partial=0.02), and Q-CHAT (β=-14.86, 95% CI=-28.45-(−1.26), p<0.05, R^2^ full=0.13, R^2^ partial=0.03). Further, network strength within the somatosensory and auditory cluster was associated with language composite (β=18.94, 95% CI=1.39-36.50, p<0.05, R^2^ full=0.10, R^2^ partial=0.02) and expressive language sub-scale (β=3.97, 95% CI=0.97-6.96, p<0.01, R^2^ full=0.07, R^2^ partial=0.04) (Figure 6).

**Figure 6.**
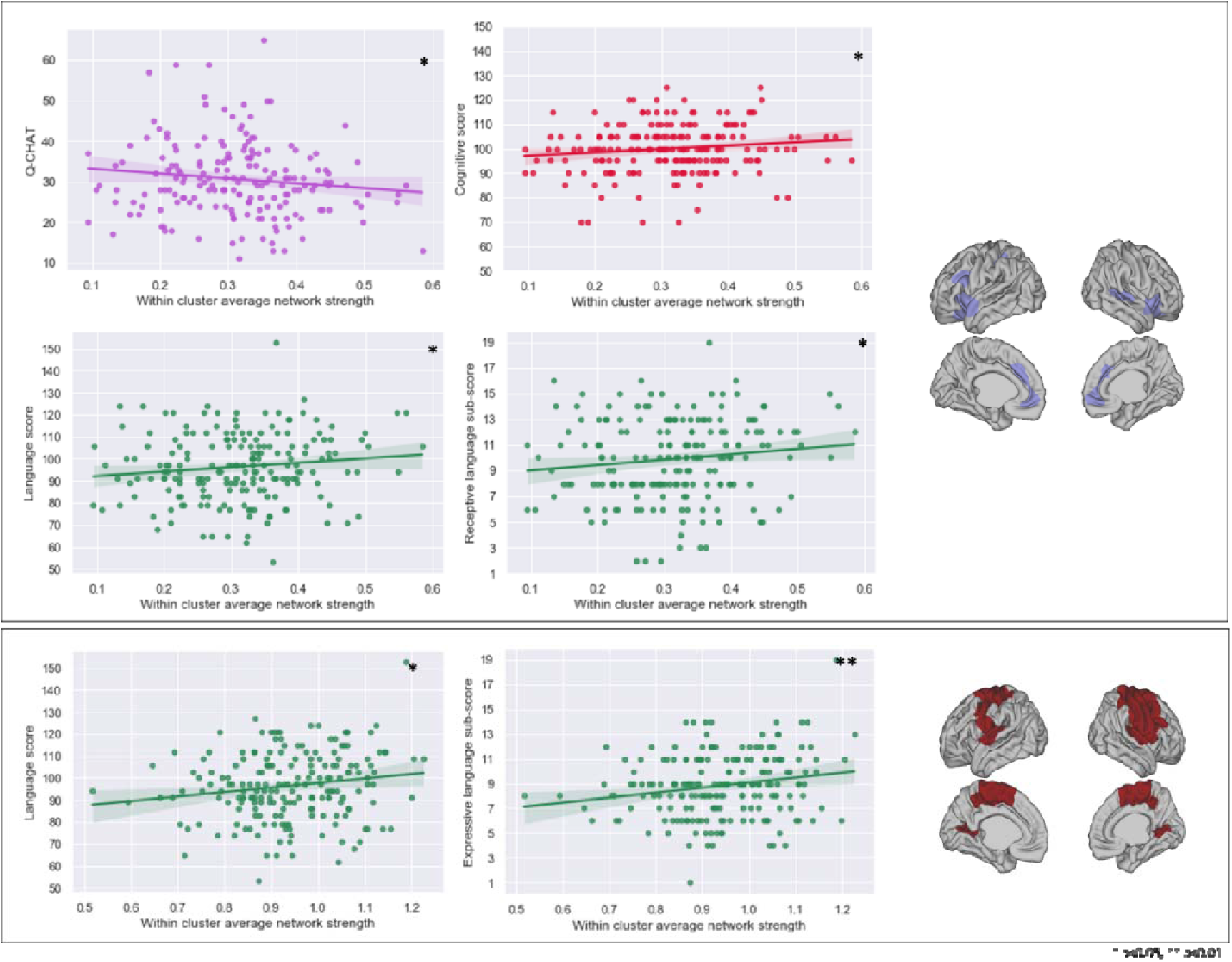
Scatterplots of significant associations between within-cluster average network strength and developmental outcomes. Plots of significant associations between within-cluster average MSN strength and social-emotional, language and cognitive measures at 18 months. Top: insular & medial frontal cluster, bottom: somatosensory & auditory cluster.

## 4. Discussion

In this study, we found that neonatal MSNs can successfully predict social-emotional behaviours at 18 months in a large group of healthy term-born babies. We show that the pattern of cortical maturation at birth already captures variability in infant development. This association between brain structure at birth and infant developmental outcomes was not limited to information held in individual edges of the network but was also evident in summary measures of network strength over the entire cortex, specifically in somatosensory-auditory, insular and medial frontal areas. On a regional level, the predictive networks were complex and widespread across the cortex, albeit suggesting specific important involvement of primary and posterior cortical regions. No consistent or significant results were found for the cognitive or motor measures.

Both language and social-emotional development result from a combination of fetal and postnatal brain programming, together with in- and ex-utero experiences. However exactly how these factors come about and interact to create these complex behaviours is still under investigation. Our results indicate that some of the neural foundations crucial for developmental capacities in infanthood, observed through the pattern of structural covariance, originate during the fetal period and are already present at birth. Social-emotional abilities were predicted by the individual edges comprising the network and associated with the average whole-network strength (Figures 2,5) implying this relationship is observed both at the micro and macro scale properties of the network.

Specifically, as a predictive network, neonatal MSNs reflect that optimal brain functioning is reliant on a delicate equilibrium of positive and negative structural covariance, where the skeleton for future infant development is established long before overt behaviours could be assessed. Our results suggest that within functional cortical divisions (i.e., insular & medial frontal and somatosensory & auditory clusters), greater structural similarity at birth is associated with better language and social-emotional skills in infanthood. This was also observed on a whole-brain level. This is in agreement with the hypothesized basis of structural covariance in the brain, whereby regions participating in similar functions show comparable structural signatures, implying joint developmental trajectories.

Predictive networks included 1%-3% of edges, consistent with other studies utilizing CPM (Cai et al. 2020; Rosenberg et al. 2016; Suo et al. 2020; Yip et al. 2019). The resulting predictive networks for social-emotional features encompassed the entire cortex, with predictive edges observed both within functional clusters and between functional clusters revealing both local and distributed networks (Figures 3-4). The predictive ability (correlation between predicted and observed values) was modest. Variance explained by whole-network strength ranged between 2%-4%, similar to previous estimates of the contribution of MRI features in explaining cognitive development in a mixed sample of term and preterm neonates (Girault et al. 2019a), but lower than estimates reported in preterm infants only (Ball et al. 2015). Combining with whole brain connectivity measures (e.g., fMRI) may provide a pathway to increasing predictive values. This reinforces the importance of other postnatal factors such as development, experience and environment (Iso et al. 2007; Koo et al. 2003) and variability in the stage and recording of cognitive development at 18 months.

The large variability in brain phenotypes during early development was demonstrated with the network predicting language abilities: While a significant predictive network emerged using LOOCV, it could not be replicated using a 10-fold cross-validation. In this case, a removal of only 10% of the subjects diminished the predictive capacity of the model. This illustrates the need for large cohorts in imaging studies of normative development in infanthood, which are now feasible through endeavours such as the dHCP. Although not robust enough, the language network revealed interesting patterns: There was some overlap between predictive networks for social-emotional and language scores. This is not surprising as these measures are conceptually related (i.e., language is critical for social communication) and correlated (*r*=-0.52). It is not only that producing and understanding speech is critical to communicating with others, but it is also postulated that social interaction and social learning experiences are crucial for the proper development of language skills (Kuhl, 2007). The overlap in the predictive networks of Q-CHAT and language scores reached a maximum of 26%, suggesting that while these scores are correlated, they also represent additional behavioural phenotypes and mechanisms. The robustness of Q-CHAT predictions but not of language skills alone may imply that utilizing a measure such as the Q-CHAT, which combines additional behaviours together with communication and language indices, may improve our ability to predict infant outcomes from brain data.

A large Australian study (Reilly et al. 2007) found that risk factors for developmental language delays such as gender, prematurity and birth weight, birth order, socioeconomic status, maternal mental health and education, and family history of language difficulties explain a small amount of variability in language abilities at age two (up to 7% of variance). Moreover, the authors found that the most predictive factor of language performance at 12 months is language performance at 8 months and that performance at 24 months is best predicted by abilities at 12 months (Reilly et al. 2006, 2007), thus indicating early trait stability. These findings support our observations that the brain basis for early capabilities begins to shape in very early life, supporting a significant role for genetic and intrauterine factors that are further influenced over post-natal development. Moreover, this shows how predicting developmental outcomes from known risk factors has very little power.

Our results point to specific involvement of occipital-parietal, somatosensory-auditory and insular-medial frontal regions. This resonates with established trajectories of brain development, whereby synapses, dendritic growth and myelination are first established in primary cortical regions (Huttenlocher & Dabholkar, 1997; Kinney et al. 1998), with cortical grey matter following a lower- to higher-order regional developmental (Gogtay et al. 2004). All of the above-mentioned regions are required for language, communication and social behaviours (Demonet et al. 2005; Porcelli et al. 2019). These larger cortical divisions include specific brain regions traditionally associated with language, such as Broca’s area (within the insular and medial/inferior frontal cluster) and the supramarginal gyrus (as part of the occipital and parietal cluster), which has been implicated in studies of toddlers with developmental language disorders using MRI (Morgan et al. 2016). Moreover, the contribution of the cortical primary motor areas complements the association between motor skills and language in both ASC and the general population (Bedford et al. 2016; Gonzalez et al. 2019), whereby better motor development is related to better language development, revealing a close relationship between these seemingly ‘independent’ domains.

Problems with social functioning and communication are associated with a variety of neurological and psychiatric conditions, most notably ASC. Research on adolescents and adults suggests that social behaviour is reliant on the proper structure and function of stand-alone brain regions, as well as of brain-wide networks (e.g., ‘amygdala network’, ‘empathy network’). Structures identified include but are not limited to the temporo-parietal junction, prefrontal cortex, superior temporal gyrus, and amygdala (Kennedy & Adolphs, 2012). From a network perspective, in young children with ASC, grey matter covariance showed patterns of decreased connectivity in the cortex, that also predicted communication scores (He et al. 2021). Nodal efficiency of tractography networks in infants who were later diagnosed with ASC was also found to be reduced in primary somatosensory, auditory and language areas (Lewis et al. 2017), where changes were detected as early as six months and related to the level of autistic symptoms at 24 months.

In previous work with smaller sample size and the inclusion of both term and preterm neonates, cortical FA at birth was found to significantly predict cognitive and language scores at two years using support vector regression (Ouyang et al. 2020), with language-related features including the inferior frontal gyrus, insula and post-central gyrus, regions that were also identified in this current work. A longitudinal study of 33 term-born infants examined the relationship between deformation-based surface distance at neonatal timepoint and the Bayley-III scores at four different time points in the first two years of life (Spann et al. 2014). The authors found significant associations between neonatal volume changes and motor and cognitive scores at 6, 12, 18 and 24 months, an association we were not able to replicate in this current work. In that study, associations with language scores were also found across all time points, highlighting the cingulate and posterior parietal areas. Interestingly, at 18 months this relationship was weaker compared to performance at 12 or 24 months.

The ontogeny of different cortical regions, and of different behavioural skills is distinct. Grossly speaking, the first cortical regions to mature are the primary areas of motor, sensory, visual and auditory functions, followed by their associative regions, and lastly maturing are the higher-order (frontal) regions (Gogtay et al. 2004). This is the case for behavioural development as well; the infant will first establish visual, somatosensory and auditory functions, before engaging in complex activities requiring language and cognitive functions. Therefore, in the immediate postnatal period, sensory, primary regions may have reached a maturational stage where enough variance is presented to detect a linear relationship with outcome. On the other hand, other regions may have not reached maturity at this point. Moreover, as the development of motor, language, cognitive, and social-emotional abilities emerge and refine at different timepoints and timescales, at 18 months, the level of expertise in these skills is different. This may indicate that certain relationships are only apparent when pairing different ages at scan and outcome and is therefore a substantial challenge when designing studies exploring early development. This was demonstrated by Hazlett et al. 2017, reporting that differences in brain volume in infants with ASC could be detected at 24 months, but not before that. Lack of results for the motor domain may relate to a narrower spread of performance in motor scores: As can be observed from Figure 5, the variability in motor scores, especially in the composite motor scores, is smaller compared to other outcome measures and could influence the ability to detect any meaningful differences.

### 4.1 Limitations

CPM has its own limitations as outlined in (Shen et al. 2017), for example, modelling only a linear relationship between variables. However, it does provide a clear data-driven framework for the implementation and interpretation of behaviour prediction models based on connectivity. We were not able to perform out-of-sample cross-validation for prediction results, only a within-sample validation and as such it remains to be confirmed whether the results generalise to a different neonatal sample. As a limitation we should also highlight that the brain-behaviour associations reported here exclude subcortical structures and the cerebellum as structural features. In this work we focused on surface-based measurements as developmental indices and therefore our analysis was limited to the cortex.

Future work should examine the contribution of subcortical regions and the cerebellum, as they are likely to also be implicated in the development of social-emotional, cognitive, language and motor abilities. Moreover, the cortical parcellation for generating the networks’ nodes was based on 150 regions. While we are aware network analysis is highly reliant on parcellation scheme, we have shown in a previous paper (see Supplementary Figure 1 in Fenchel et al. (2020)), that MSNs could be replicated using random partitions of different sizes. Given that the neonatal brain is also substantially smaller, in this current work we chose the n=150 parcellation as a middle ground between granularity and interpretability, while not sampling very small patches.

When interpreting scores from developmental assessments, such as the ones used in this study, one should understand their nature: One assessment at 18 months is only a transient screenshot of infant development and is not necessarily indicative of future difficulties at the individual level. Both language delays and the appearance of social-emotional difficulties at an early age do not inevitably mean the continuation of language difficulties or a later autism diagnosis. In addition, these features or delays might not be evident at 18 months but only emerge at a later age. Therefore, any interpretation of infant assessments in the context of future functionality should be done with caution and preferably include follow-up at preschool and school-age (Duff et al. 2015; Fountain et al. 2012; Waizbard-Bartov et al. 2021).

### 4.2 Conclusions

In this work, we showed that multi-feature multi-modal cortical similarity at birth represented by MSNs are predictive of social-emotional abilities in a large group of infants. Cortical regions involved were widespread, with predictive features including connectivity within and across functional cortical domains. Earlier developing cortical regions seemed to be specifically important in this context. These results support the use of neonatal cortical profiles for means of early detection and support of developmental difficulties.

## Acknowledgements

We would like to thank the infants and parents who contributed their time to this research and the research team at the Newborn Imaging Centre at Evelina London. We are also grateful for comments made by anonymous reviewers.

