## Supplementary Material for "Neonatal multi-modal cortical profiles predict 18-month developmental outcomes"

Supplementary Table 1. Pearson’s correlation between the seven Bayley Scales of Infant and Toddler Development (Bayley-III) items and Q-CHAT

|  | Language | EL | RL | Motor | FM | GM | Q-CHAT |
| --- | --- | --- | --- | --- | --- | --- | --- |
| Cognitive | 0.53*** | 0.46*** | 0.51*** | 0.47*** | 0.45*** | 0.26*** | -0.36*** |
| Language |  | 0.90*** | 0.93*** | 0.47*** | 0.39*** | 0.33*** | -0.52*** |
| EL |  |  | 0.67*** | 0.41*** | 0.33*** | 0.30** | -0.41*** |
| RL |  |  |  | 0.45*** | 0.38*** | 0.31*** | -0.52*** |
| Motor |  |  |  |  | 0.81*** | 0.74*** | -0.28*** |
| FM |  |  |  |  |  | 0.20** | -0.22** |
| GM |  |  |  |  |  |  | -0.21** |

** p<0.05, ** p<0.01; SE-Standard error, EL- expressive language, RL- receptive language, FM- fine motor, GM- gross motor, Q-CHAT- Quantitative Checklist for Autism in Toddlers.

Supplementary Figure 1. Prediction of language scores from neonatal MSNs using CPM.

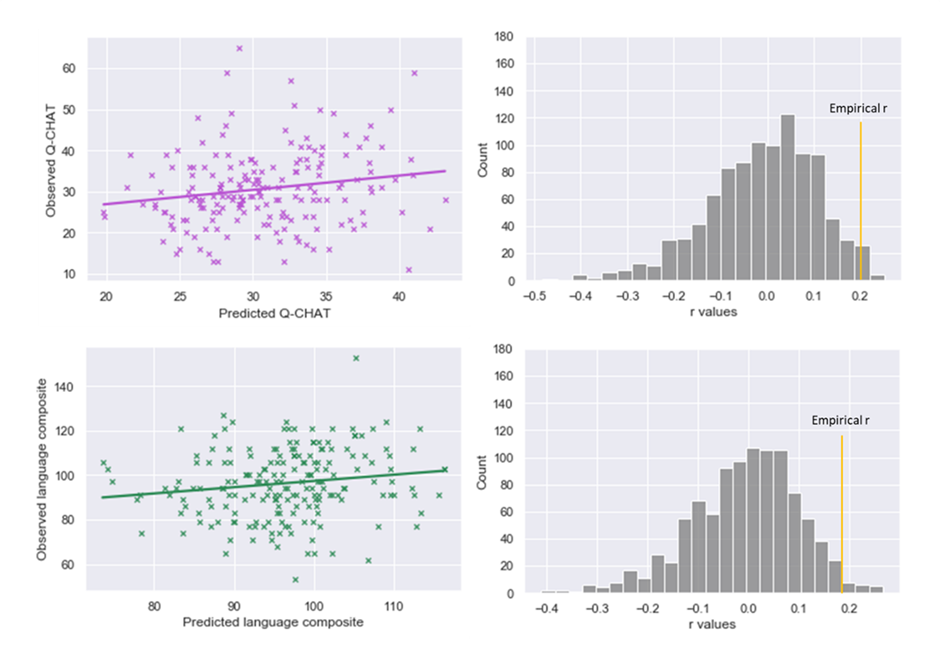

Plots of significant correlation between predicted and observed language scores (left) and results of null r values with permutation testing (right) using Connectome-based Predictive Modeling (CPM).

Supplementary Figure 2. Proportion of between-cluster edges involved in language networks.

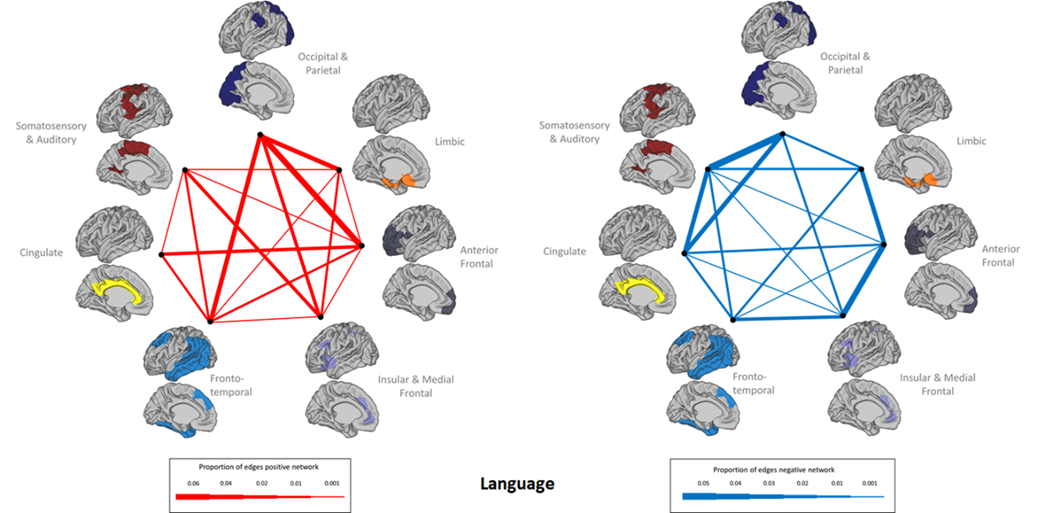

Proportion of edges included in language prediction model connecting nodes between clusters. On the left the positive network is shown and, on the right, the negative network.
